# Quantification of cellular phototoxicity of organelle stains by the dynamics of microtubule polymerization

**DOI:** 10.1101/2024.01.17.576021

**Authors:** Shivam Mahapatra, Seohee Ma, Bin Dong, Chi Zhang

## Abstract

Being able to quantify the phototoxicity of dyes and drugs in live cells allows biologists to better understand cell responses to exogenous stimuli during imaging. This capability further helps to design fluorescent labels with lower phototoxicity and drugs with better efficacy. Conventional ways to evaluate cellular phototoxicity rely on late-stage measurements of individual or different populations of cells. Here, we developed a quantitative method using intracellular microtubule polymerization as a rapid and sensitive marker to quantify early-stage phototoxicity. Implementing this method, we assessed the photosensitization induced by organelle dyes illuminated with different excitation wavelengths. Notably, fluorescent markers targeting mitochondria, nuclei, and endoplasmic reticulum exhibited diverse levels of phototoxicity. Furthermore, leveraging a real-time precision opto-control technology allowed us to evaluate the synergistic effect of light and dyes on specific organelles. Studies in hypoxia revealed enhanced phototoxicity of Mito-Tracker Red CMXRos that is not correlated with the generation of reactive oxygen species but a different deleterious pathway in low oxygen conditions.

**Teaser:** Microtubule dynamics in live cells allow quantification of cellular phototoxicity of fluorescent dyes in various conditions

## Introduction

When applying optical microscopy to imaging live biological samples, minimizing disruptions to biological functions is critical. The use of lasers to excite the fluorescent molecules may result in undesirable phototoxic effects (*1–4*). For instance, the excited fluorophore might transport energy to neighboring molecules, inducing physicochemical changes via photosensitization. The excited fluorophores might also degrade into more toxic configurations that impair cellular functions. In a nutshell, phototoxicity can modify the biological functions within a live sample, leading to potential misinterpretation of acquired results and unwanted sample alterations. Hence, the assessment and reduction of phototoxicity represent a primary objective in optical microscopy, particularly when employing laser scanning methods due to the concentrated laser intensity at the focal point.

Most of the existing methods to quantify cellular toxicity rely on measuring collective cell responses at later stages. For example, the phototoxicity of dyes to live cells is often quantified using cell viability assays such as 3T3 neutral red uptake, MTT, alamarBlue, or lactate dehydrogenase (LDH) release after illumination (*5–8*). Apoptosis and necrosis assays can also be utilized (*9*). These assays primarily capture late-stage cellular responses and lack sensitivity at the individual cell level. They also fall short in measuring the phototoxicity arising from optical microscopy since only a small fraction of cells are illuminated during imaging. Alternatively, timelapse microscopy offers a means to observe shifts in dynamic cellular processes (*1, 4*). Alterations in cell morphology and mobility serve as characteristics for assessing phototoxicity. Cellular stress, for example, can induce changes like cell shrinkage, mitotic delay, vacuole formation, or membrane blebbing, significantly altering cell morphology (*10–12*). Furthermore, the mobility of cells can also be a readout (*13*). Despite having a single-cell resolution, these readouts generally focus on late-stage cellular responses. Intracellular dynamics of organelles, regulated by molecular motors and fueled by metabolites, can respond to perturbations within an hour (*14, 15*). Intracellular fluorescence signals can provide faster measurements of cell responses to stimuli. For instance, the sensors responding to intracellular reactive oxygen species (ROS) offer sensitivity to quantify ROS during regular cellular processes or under stress (*16, 17*). The ROS indicators also aid in evaluating phototoxic effects since photosensitization can lead to elevated ROS generation (*3, 5, 18*). However, the introduction of exogenous ROS indicators requires uptake time and would perturb cell functions (*19*).

In this study, we first report a sensitive and quantitative method to measure early-stage cellular phototoxicity based on fluorescent protein signals tracing the dynamics of microtubule polymerization. The visualization of real-time microtubule polymerization dynamics is achieved by co-expressing enhanced green fluorescence protein (EGFP) with end-binding protein 3 (EB3) in HeLa cells. The quantification of EB3 comets provides a sensitive assessment of cellular alterations. This approach offers fast response time, single-cell sensitivity, and avoids perturbation to biofunctions. Applying this method to commonly used organelle labels, we quantify the induced phototoxicities, which are found to be both chemical-dependent and excitation wavelength-dependent. Furthermore, a real-time precision opto-control (RPOC) technology is applied to compare cell responses when illuminating labeled and unlabeled areas. The distinct differences in cell responses validate the synergistic effect of the excitation light and dyes interacting at selected organelles. Moreover, our investigation into the impact of oxygen levels on phototoxicity revealed that Mito-Tracker induces increased phototoxic effects in hypoxic conditions, which operate through a mechanism distinct from ROS. The methodologies and findings presented in this study offer an approach for early-stage assessment of cellular responses to photoperturbation, providing novel insights into phototoxicity induced by diverse fluorescent labels targeting various organelles.

## Results

### Quantification of microtubule polymerization dynamics using EB3 comets

Phototoxicity of a chemical is defined as the derived toxicity of the chemical upon exposure to light (*4*). In other words, the interaction between the chemical and the absorbed light produces toxic intermediates that disturb cellular homeostasis. Understandably, the impact is most pronounced when the excitation laser closely matches the wavelength at which the chemical exhibits its highest light absorption. Conventional phototoxicity assays based on ensemble cell analysis fail to detect early-stage cell responses and lack single-cell sensitivity. Light microscopy monitoring of phototoxicity typically rely on significant cellular morphological changes at the late-stage of perturbation. The use of exogenous fluorescent sensors can induce additional functional perturbations to cells, thus complicating the analysis of phototoxicity. We developed a quantitative method using microtubule polymerization dynamics as a marker for sensitive and non-perturbative analysis of phototoxicity in live cells.

To visualize the dynamics of the microtubules, we use HeLa cells with EGFP transfected and co-expressed with EB3, a dynamic protein that preferentially binds to the plus end of the microtubules (*20*). EB3 regulates the dynamics of the microtubule growth via binding to the ‘guanosine triphosphate (GTP) cap’ which contains GTP-bound tubulin that stabilizes its growth (*21, 22*). The binding dynamics of EB3, therefore, reflect the presence of GTP-bound tubulin dimers and GTP levels in cells (*22*). Cellular GTP level, due to its direct relationship with the Krebs cycle, mitochondria functions, and GTPase activity, is a sensitive indicator of the overall energy status and metabolism of the cell (*23, 24*). Therefore, monitoring the highly dynamic EB3-binding process provides a quick and reliable quantification of changes in cellular energetic states that have fast responses to phototoxicity. We discovered that quantification of EB3 comets provides a rapid and sensitive measure of phototoxicity induced by various factors in live cells. Moreover, the EGFP fluorophore, enclosed within a protective polypeptide envelope, remains non-phototoxic during excitation.

A time-lapse video of EB3 dynamics in HeLa cells is shown in **Supplementary Video 1**. The EB3 comets typically grow at a speed of about 0.2-0.4 µm per second and shrink even faster (*25*). Phototoxic perturbations of live cells change the energy flux of cells and consequently alter the length and intensity of the EB3 comets. To measure the change in comet length, we developed a quantitative method as illustrated in **Figure 1A**. First, a time-lapse fluorescence image stack is acquired. Next, a denoising method is applied to enhance the contrast of EB3 comets (*26*). The first and last denoised output images undergo processing with a Gaussian-blur filter, followed by subtraction from the original image. This process effectively minimizes the background inhomogeneity and highlights the intensity variations associated with the comets. Then, an intensity threshold is used to isolate comets for quantification, followed by "Particle analysis” to calculate the circularity of each particle. The circularity is defined as

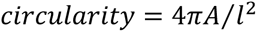

**Figure 1.**
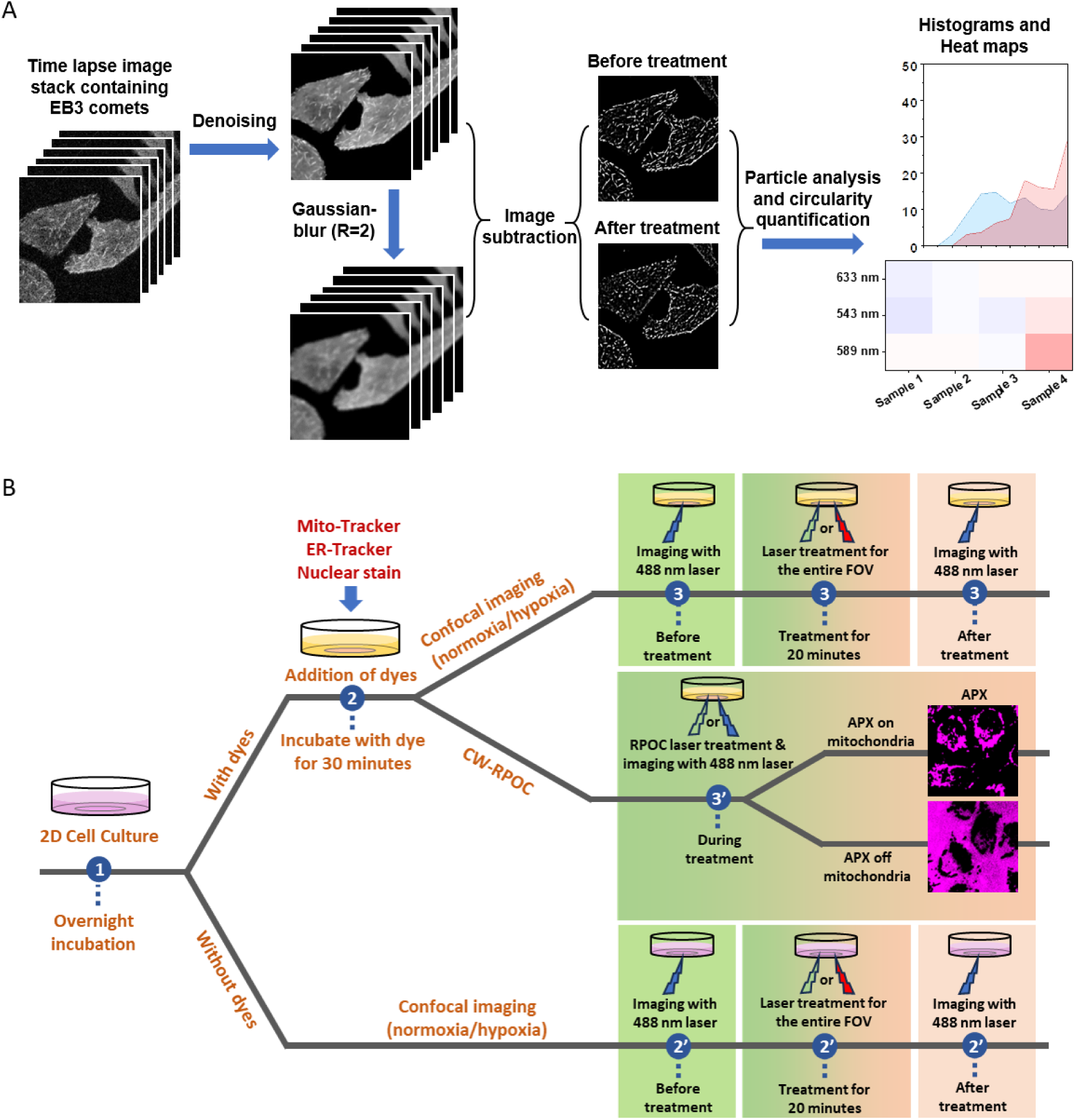
Assessing the cellular phototoxicity caused by various fluorescent organelle dyes and laser wavelengths in live cells. (A) The procedure for quantifying phototoxicity using EB3 comets and microtubule polymerization dynamics. (B) Experimental frameworks for analyzing phototoxicity of fluorescent organelle dyes when excited by different laser wavelengths. The continuous wave real-time precision opto-control (CW-RPOC) technology permits the examination of the synergistic effects of lasers and dyes on specific organelles.

where *A* is the total area of the particle, and *l* is the particle perimeter. A circularity value of 1 signifies a perfect circle, while a decrease in the circularity value indicates increased elongation of the particle. Longer EB3 comets yield smaller circularity values compared to shorter ones. The various circularity values of EB3 comets from each image are graphed in a statistical histogram. Furthermore, the percentage of particles exhibiting a circularity below 0.6 is quantified to generate a heat map. Note that this method is generally applicable across all confocal microscopes and is independent of the image size, provided individual comets are detectable and exhibit similar lengths within the control group. However, when employing different imaging systems, the circularity values may vary slightly, necessitating results from control groups as references.

### Evaluation of the phototoxicity of organelle markers

The study design of investigating the phototoxic effects of commonly used fluorescent organelle dyes is illustrated in **Figure 1B**. Optical treatment at different wavelengths is compared for cells labeled with various organelle dyes and their label-free counterparts in the same microenvironment. To determine whether phototoxicity arises due to light-dye synergistic interactions at specific organelles, we employ a recently developed RPOC technology. RPOC activates or deactivates lasers solely at desired molecular targets while simultaneously monitoring cell responses (*27–29*). Within the RPOC framework, an active pixel (APX) is defined as the pixel at which the action laser is activated. In this study, we label mitochondria with Mito-Tracker and compare phototoxicity when the laser APXs are selected on and deselected from mitochondria.

First, we treated cells with the nucleus dye SYTOX Orange Nucleic Acid Stain, which exhibits a maximum absorption wavelength of 547 nm and an emission wavelength of 570 nm. The phototoxic impact of this dye is very significant, leading to the disappearance of EB3 comets within 25 seconds, even in the absence of the optimal excitation laser. As shown in **Figure S1**, continuous illumination of cells solely with the 488 nm laser that excites the EGFP-EB3 signals, results in a notable decline in microtubule polymerization dynamics. Expanding the field of view (FOV) reaffirms these alterations as light-induced, since the unilluminated area still exhibits unaffected EB3 dynamics. When subjected to the illumination of 543 nm laser that matches the excitation wavelength of the nucleus dye, the EB3 comets disappear within a single frame, indicating a rapid cessation of microtubule polymerization.

Next, the phototoxicity of ER-Tracker Red (Ex 587/Em 615) is studied. In comparison to the nucleus dye, this particular dye exhibits significantly reduced phototoxic effects. Illuminating unlabeled cells with 543 nm and 633 nm lasers for 20 minutes shows undetectable changes in EB3 comets in both imaging and quantitative circularity analysis (**Figure 2A**). When the ER-Tracker is applied, only a slight change in EB3 comet circularity is detected upon 20 minutes of illumination with the 543 nm laser (**Figure 2B**). Given that the excitation peak of the ER-Tracker is approximately 587 nm which is beyond the laser options of our commercial confocal fluorescence microscope, we treat the cells using a 589 nm laser in a lab-built confocal system and monitored the EB3 signal changes. Despite unchanged microtubule dynamics observed without the dye (**Figure 2C**), a significant decrease of EB3 comets is discovered after about only 150 s of 589 nm laser illumination with the ER-Tracker (**Figure 2D**). Such a change is evident in both fluorescent images and comet circularity histograms. **Figure 2E** displays the absorption spectrum of ER-Tracker alongside the three wavelengths utilized in this investigation. The 589 nm excitation induces the most pronounced phototoxic effect on HeLa cells due to its strong absorption by the ER-Tracker. To quantitatively assess phototoxicity under varying conditions, a heatmap plotting the percentage of EB3 comets with circularity below 0.6 is generated (**Figure 2F**). Lower values on the heatmap signify increased phototoxicity.

**Figure 2.**
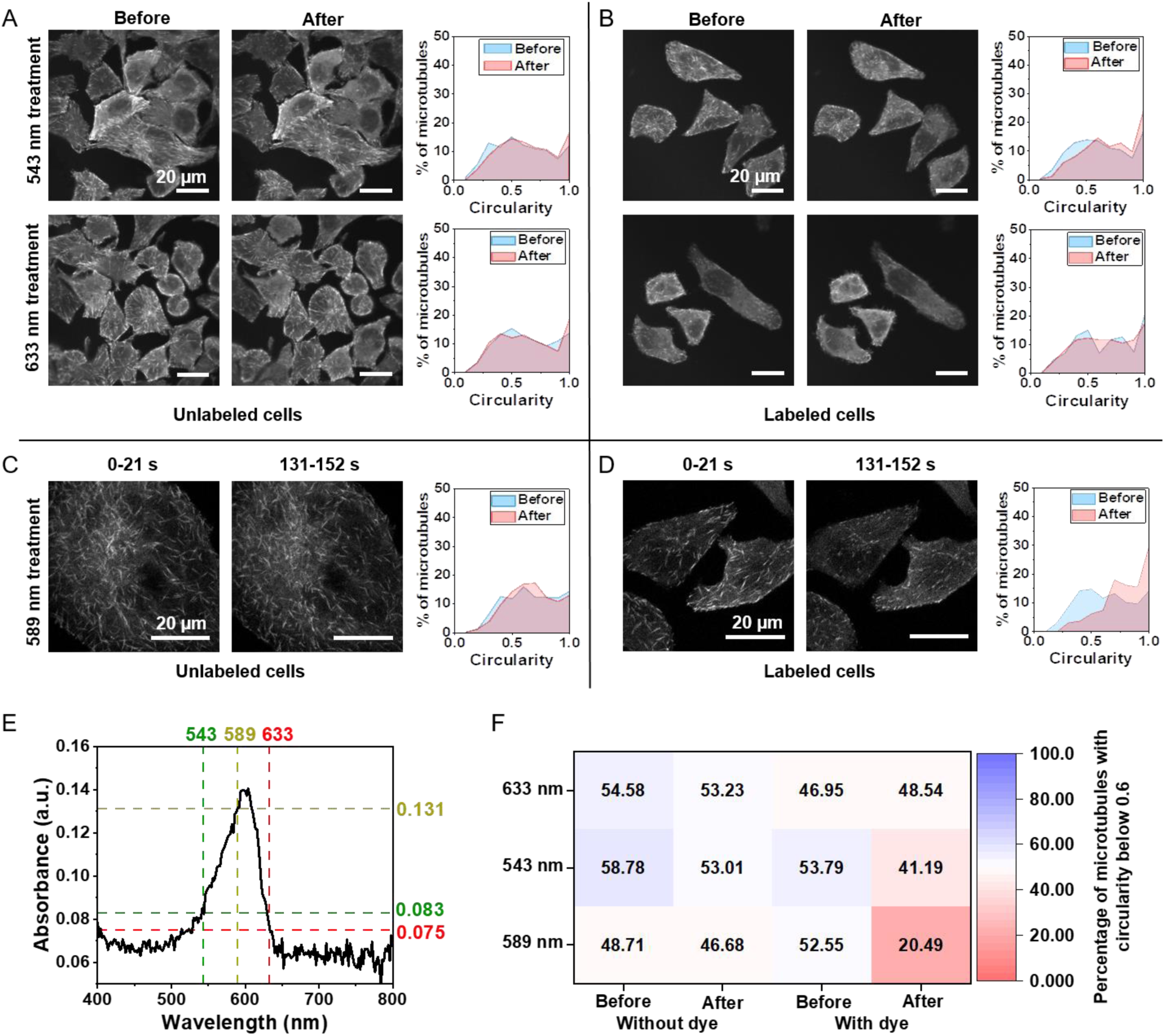
Quantification of the cellular phototoxicity induced by ER-Tracker under the excitation of different laser wavelengths. (A) Fluorescent images of EB3-EGFP signals excited by 10 µW 488 nm laser and circularities of EB3 comets from HeLa cells before and after illumination by 543 nm and 633 nm lasers for 20 minutes via laser scanning. (B) Similar to panel A, where the cells are stained with 1 µM ER-Tracker (587/615 nm). (C) Fluorescent images of EB3-EGFP signals and circularities of EB3 comets from HeLa cells illuminated by 589 nm laser for 152 s using a lab-built confocal fluorescence microscope. A 20 µW 473 nm laser is also scanned with the treatment laser to excite the EGFP signals. (D) Similar to panel C, where the cells are labeled with 1 µM ER-Tracker (587/615 nm). (E) The absorption spectrum of ER-Tracker (587/615 nm). (F) A heatmap showing the percentage of EB3 comets with circularities below 0.6 for each treatment condition.

A similar investigation is conducted for Mito-Tracker Red CMXRos (Ex 579/Em 599). The absorption spectrum of Mito-Tracker Red, displayed in **Figure S2**, exhibits a broad absorption peak with the maximum absorption around 590 nm, similar to the ER-Tracker Red. In the case of unlabeled cells, no evidence of phototoxicity emerges when either the 543 nm or 633 nm laser is illuminated for 20 minutes (**Figure 3A**). However, upon application of Mito-Tracker to HeLa cells, the 543 nm laser eliminates all EB3 comets after treatment while the 633 nm laser shows no discernible effect (**Figure 3B**). Notably, prominent membrane blebbing and vacuole formation are evident in most cells in the FOV after the 543 nm laser treatment. For the cells at the edges of the FOV, some observable EB3 comets remain, as only a portion of the cells is illuminated. It’s important to note that the reduction in EB3 comets induced by phototoxicity typically amplifies the overall fluorescent intensity of the cytosol. This phenomenon occurs because depolymerized EB3 molecules are released from polymerizing microtubules into the cytosol. To better match the optimal excitation wavelength, a 589 nm laser from a lab-built confocal microscopy is employed for excitation. Although the 589 nm laser does not affect microtubule polymerization in the absence of MitoTracker, the cells labeled with MitoTracker exhibit the disappearance of EB3 comets within 150 seconds upon interaction with the 589 nm laser, as depicted in **Figure 3D**. Because the EB3 comets within MitoTracker labeled cells are entirely eradicated following interactions with 543 nm and 589 nm lasers, as depicted in **Figure 3**, the circularity analysis is not conducted. Literature also provide evidence of the photosensitizing action of Mito-Tracker Red CMXRos. Upon exposure to light, this dye induces damage to the inner membrane of the mitochondria, leading to mitochondrial swelling and cell death (*30*). Moreover, it causes a drastic change in the mitochondria movement in *Arabidopsis thaliana* protoplasts (*31*).

**Figure 3.**
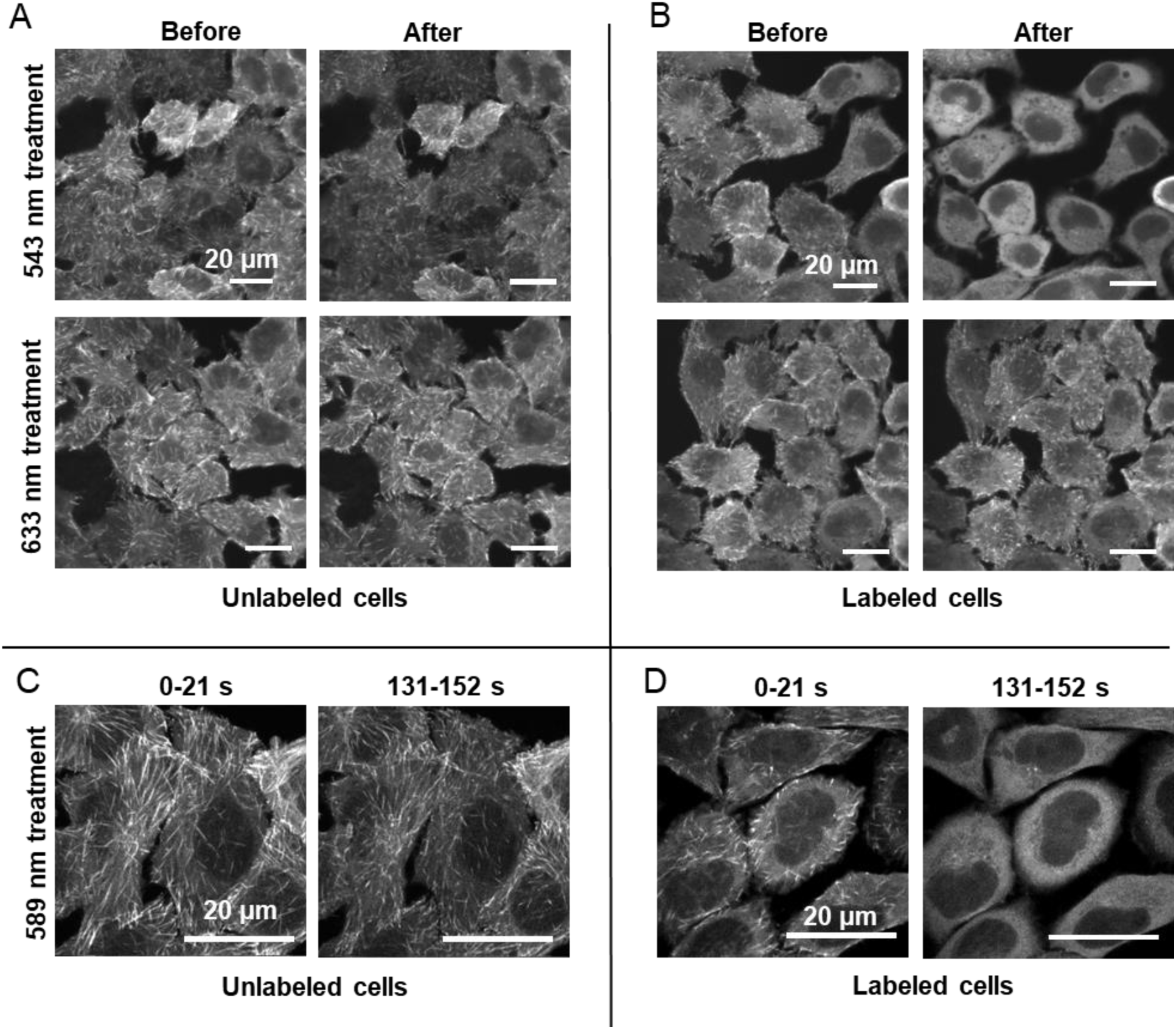
Cellular phototoxicity of Mito-Tracker under the excitation of different laser wavelengths. (A) Fluorescent images of EB3-EGFP signals from HeLa cells excited by 10 µW 488 nm laser before and after illumination by 543 nm and 633 nm lasers for 20 minutes via laser scanning. (B) Similar to panel A, where the cells are labeled using 200 nM Mito-Tracker Red CMXRos. The 543 nm laser excitation induces strong phototoxicity that eliminates the EB3 comets after treatment. (C) Fluorescent images of EB3-EGFP signals from HeLa cells illuminated by 589 nm laser for 152 s using a lab-built confocal fluorescence microscope. A 20 µW 473 nm laser is also scanned with the treatment laser to excite the EGFP signals. (D) Similar to panel C, where the cells are labeled using 200 nM Mito-Tracker Red CMXRos.

Overall, our collective findings demonstrate that under similar imaging conditions, the excitation of MitoTracker results in more significant phototoxic effects compared to the ER-Tracker. Assessing microtubule dynamics through EB3 comets offers a fast and quantitative way to evaluate and compare phototoxicity induced by different dyes and laser wavelengths.

### Localized photochemical perturbations

Despite our results indicating that the maximum phototoxicity occurs when the cells are labeled with the fluorescent dyes and illuminated with the corresponding excitation laser wavelength, there remains uncertainty regarding whether the synergy between the laser and dye is spatially colocalized. In other words, is the laser impacting solely the labeled organelles or other locations of the cell? Conventional imaging methods do not permit such a comparison. To address this, we developed RPOC technology that allows us to activate lasers exclusively at targets of interest (*27–29*). RPOC is based on a laser scanning microscope and a fast optoelectronic feedback system. The lasers employed for optical manipulation or perturbation are activated only at identified chemical targets during laser scanning (*27–29*). Its responsiveness and decision-making speed operate at sub-microsecond levels, thus enabling the detection of chemical targets, simultaneously inducing optical perturbation solely at these targets, and real-time monitoring of cellular responses. A schematic illustration of RPOC is shown in **Figure S3**.

We label mitochondria with Mito-Tracker and apply RPOC to select APXs only on mitochondria and excluding mitochondria. Within our RPOC setup, a 473 nm laser is employed to excite both the EB3-EGFP and MitoTracker signals and is scanned across the entire FOV. Both a 405 nm laser and a 532 nm laser are available as action lasers for precise site-specific optical perturbation. Specificially, the 532 nm laser is utilized here to optically illuminate the APXs that co-localize with Mito-Tracker signals. Simultaneously, we monitored changes in both EB3 and Mito-Tracker signals using separate fluorescence detection channels. Our findings reveal that casting the 532 nm laser on mitochondria for about 150 s reduces microtubule polymerization (**Figure 4A**). However, when the laser is directed away from Mito-Tracker-marked mitochondria, it demonstrates minimal impact on EB3 comets and microtubule polymerization (**Figure 4B**). This disparity can be better compared in the heat map displaying comet circularity below 0.6 (**Figure 4C**). In contrast to **Figure 3B**, the less pronounced disruption of microtubule polymerization in RPOC is attributed to the considerably shorter duration of laser exposure. Furthermore, compared to **Figure 3D**, the less significant changes observed are due to the mismatch between the excitation wavelength and dye’s absorption maximum.

**Figure 4.**
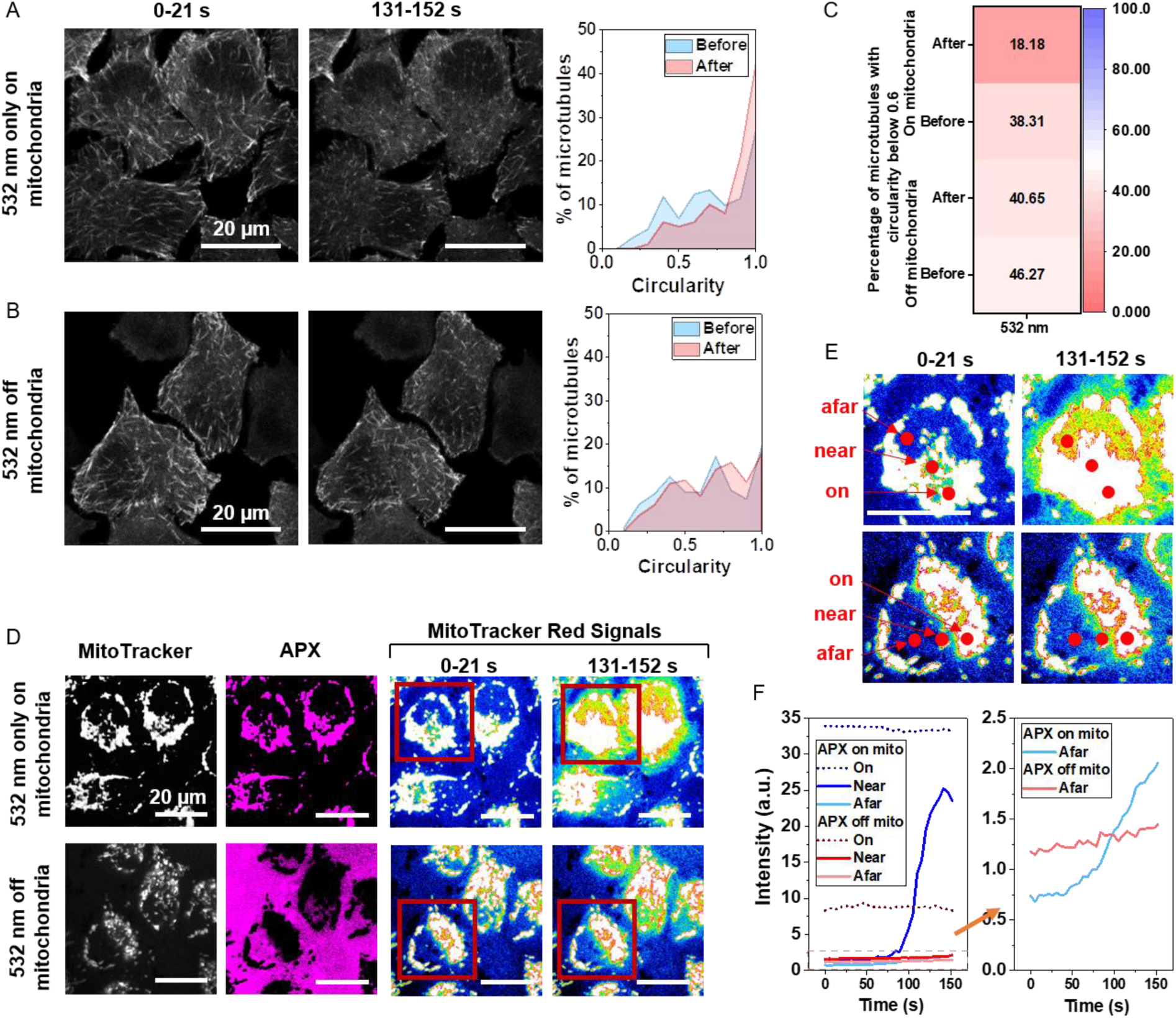
RPOC for the analysis of the synergistic impact of Mito-Tracker and lasers specifically at mitochondria. (A) Fluorescent images of EB3-EGFP signals and circularities of EB3 comets from HeLa cells illuminated by a 532 nm laser only at the mitochondria. A 20 µW 473 nm laser is also scanned with the treatment laser to excite the EGFP signals. Active pixels (APXs) are chosen to align with Mito-Tracker and the mitochondria. (B) Similar to panel A, where the APXs are deselected from Mito-Tracker and mitochondria. (C) A heatmap showing the percentage of EB3 comets with circularities below 0.6 for each treatment condition. (D) The APXs and Mito-Tracker signals before and during the treatment of 532 nm laser on and off mitochondria. During treatment, the MitoTracker signals are displayed using a 16-color scheme to better illustrate the leakage of MitoTracker into the cytosol. (E) Enlarged images for the framed areas in panel D. ‘On’ indicates on mitochondria; ‘Near’ indicates outside but proximal to mitochondria; ‘Afar’ indicates outside and distant from mitochondria. All selected locations are within the same cell. (F) MitoTracker fluorescent signals at various subcellular locations designated in panel E. The dashed range in the left panel is magnified and shown in the right panel.

In the Mito-Tracker channel, we detect a significant fluorescence signal enhancement at mitochondria when APXs are present on Mito-Tracker, contrasting with no signal increase at mitochondria when APXs exclude Mito-Tracker (**Figure 4D,E**). Furthermore, the diffusion of Mito-Tracker outside mitochondria is detected when APXs are associated with these mitochondria (**Figure 4E**). This diffusion manifests as an elevation in the Mito-Tracker signal in locations proximal or distal to the mitochondria. It significantly increases the concentration of Mito-Tracker in the vicinity of mitochondria, resulting in a cascading effect that activates APXs at those locations. Consequently, there is a substantial surge in the Mito-Tracker signals within the channel (**Figure 4E**). The leakage of Mito-Tracker into the cytosol likely stems from the phototoxicity of Mito-Tracker that disrupts the mitochondrial membrane potential (*30*). On the contrary, the dye leakage is minimal when mitochondria are excluded from APXs. In this study, RPOC is implemented under an oversampling condition, where the pixel size is approximately 100 nm, significantly smaller than the laser beam spot at the sample. The weak Mito-Tracker leaking, when the APXs and mitochondria do not overlap, is likely caused by the 532 nm laser illumination spots extending beyond the APXs solely at their boundaries or the impact of the 473 nm excitation laser scanning throughout the sample.

The RPOC study demonstrates that the phototoxic effects induced by the 532 nm laser and Mito-Tracker stem from their synergistic interactions specifically localized within the mitochondria.

### Phototoxicity of organelle markers in hypoxia

The toxicity of photosensitizers is usually linked to the generation of ROS, either through excited state energy transfer to a substrate molecule (Type I photosensitizers) or to molecular oxygen (Type II photosensitizers) (*32*). In both scenarios, the generated ROS is regarded as the primary responsible factor. To investigate the mechanisms of Mito-Tracker Red CMXRos and ER-Tracker Red, we conduct a comparative analysis of their photosensitization effects under normoxia and hypoxia conditions. The hypoxic condition is achieved by reducing the oxygen levels to 0.1% using a stage-top incubator (**Figure S4**).

In normoxia, a 20-minute exposure to a 543 nm laser significantly diminishes microtubule polymerization in cells labeled with Mito-Tracker (**Figure 5A**). Under the hypoxic condition for untreated cells, while there is an overall decrease in the intensity of EB3 comets, their presence remains largely unchanged (**Figure 5A and S5A**). This suggests that despite the reduction in intensity, hypoxia does not substantially alter microtubule polymerization. However, following treatment of Mito-Tracker-labeled cells with a 543 nm laser in hypoxic conditions, more pronounced chemical and morphological changes are observed. Both the elimination of EB3 comets and significant membrane blebbing are detected (**Figure 5A and S5A**). The overall increase in fluorescent signal in the EB3 channel indicates diffusion of Mito-Tracker and possible alteration of this dye post-interaction with the 543 nm laser. Using a fluorescent ROS sensor dichlorodihydrofluorescein diacetate (H2DCFDA), our imaging results indicate a significantly reduced ROS generation in hypoxia compared to normoxia (**Figure 5B,C, and S5B**). Therefore, the enhanced phototoxicity induced by the 543 nm laser illumination on Mito-Tracker Red in hypoxia does not appear to be associated with ROS. Given the photosensitization occurring in mitochondria, this result indicates the critical importance of mitochondrial function for cells under hypoxic conditions.

**Figure 5.**
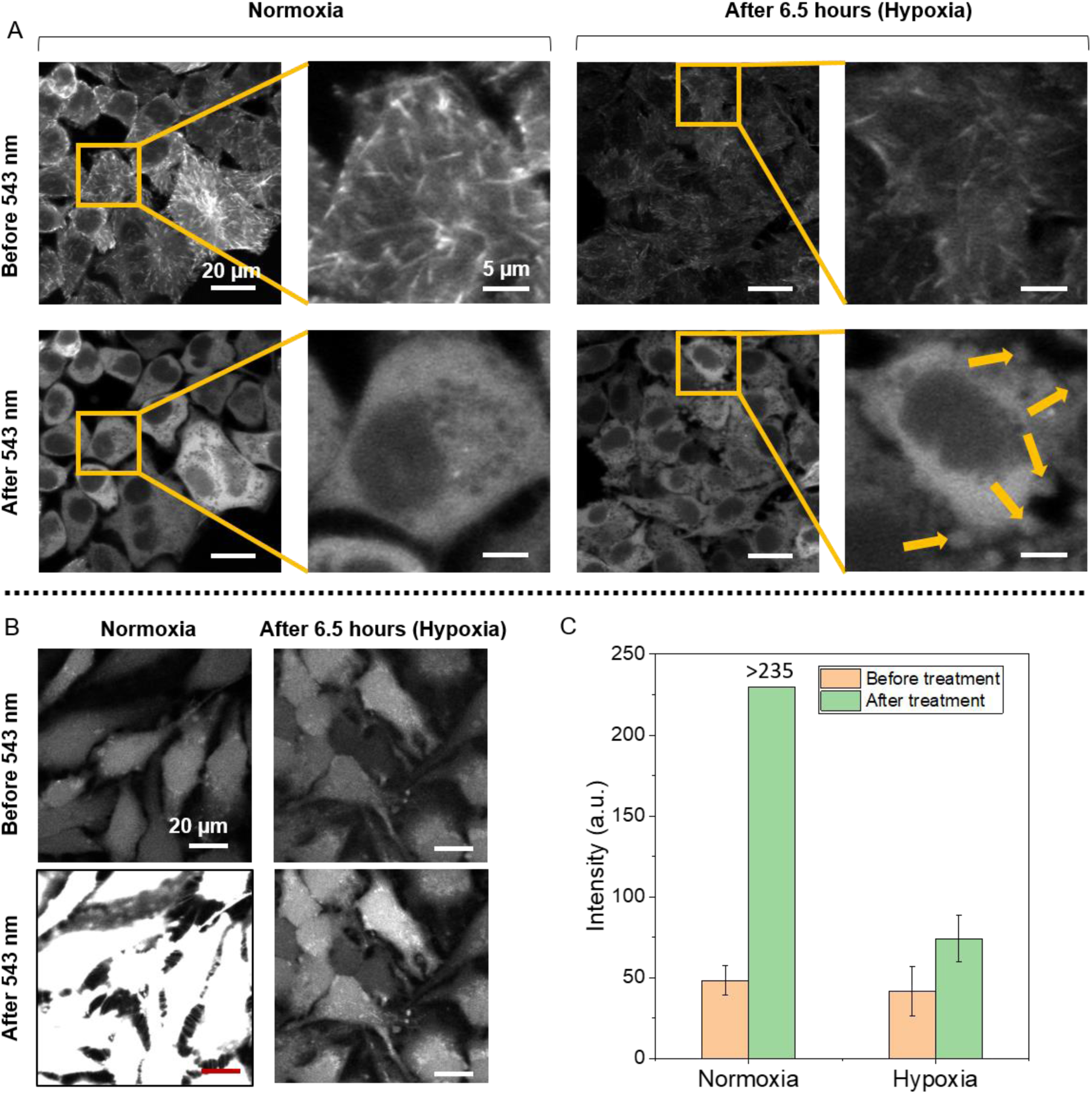
Assessment of cellular phototoxicity and ROS generation upon illuminating MitoTracker using a 543 nm laser in both normoxia and hypoxia conditions. (A) Fluorescent images of EB3 comets from HeLa cells illuminated by the 543 nm laser for 20 min in hypoxia and normoxia conditions. A 10 µW 488 nm laser is used to excite the EGFP signals. Enlarged sections within the images provide enhanced cellular details. The treatment in hypoxia induces significant membrane blebbing, as indicated by the arrows, suggesting an elevated cellular phototoxicity. (B) ROS generation induced by the MitoTracker pre and post 543 nm laser illumination in normoxic and 6.5 hr hypoxic cultured conditions. The fluorescent signals originate from H2DCFDA incubated with HeLa cells. A 10 µW 488 nm laser is used to excite the H2DCFDA signals. (C) Average H2DCFDA intensities observed in cells under various conditions in panel B.

For ER-Tracker labeled cells, a brief exposure to a 589 nm laser only slightly reduces EB3 comets and microtubule polymerization in normoxia (**Figure 6A**). In hypoxia, no notable increase in phototoxic effects is detected after 152-second treatment by the 589 nm laser (**Figure 6A and S6A**). Moreover, a longer 20-minute treatment with the 589 nm laser does not demonstrate enhanced phototoxicity in hypoxia for ER-Tracker labeled cells either (**Figure S7**). Utilizing a fluorescent ROS sensor, we identified a reduction in ROS generation in hypoxia when ER-Tracker is exposed to the 589 nm laser, indicated by a slower increase in H2DCFDA signal (**Figure 6B,C, and S6B**). Unlike the impact observed with MitoTracker treatment, the phototoxicity associated with ER-Tracker does not appear to be amplified under hypoxic conditions.

**Figure 6.**
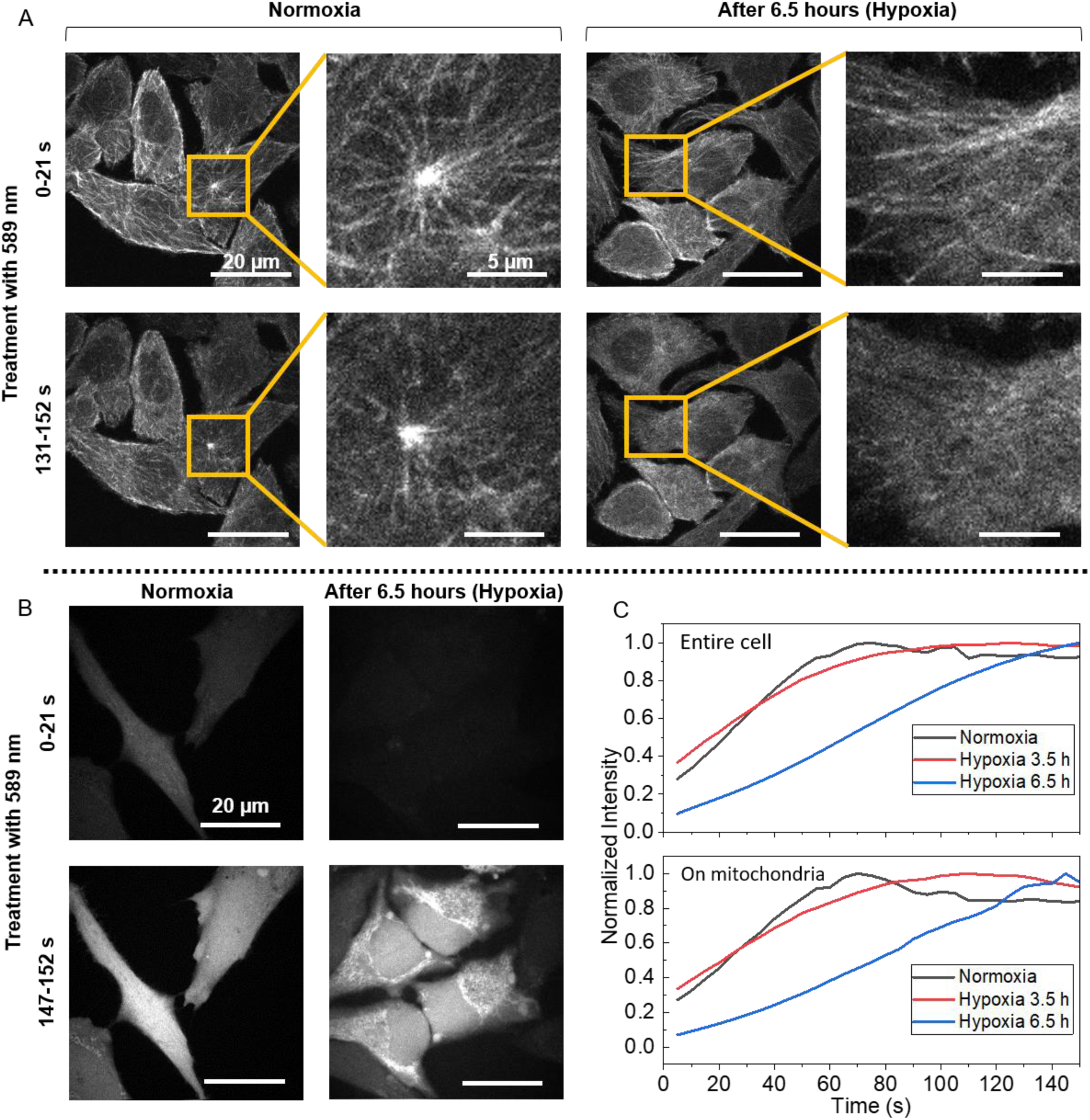
Cellular phototoxicity and ROS generation due to ER-Tracker illumination by a 589 nm laser in normoxia and hypoxia conditions. (A) Fluorescent images of EB3 comets from HeLa cells illuminated by the 589 nm laser for 152 s in hypoxia and normoxia conditions. A 20 µW 473 nm laser is scanned alongside the treatment lasers in all cases to excite the EGFP signals. Enlarged sections within the images provide enhanced cellular details. (B) ROS generation induced by the MitoTracker pre and post 589 nm laser illumination in normoxia and 6.5-hour hypoxic culture. The fluorescent signals originate from H2DCFDA incubated with HeLa cells. A 20 µW 473 nm laser is used to excite the H2DCFDA signals. (C) The alterations in H2DCFDA signals during laser treatment. Slower rises in signals suggest reduced ROS generations.

These findings highlight the diverse phototoxic effects exhibited by different organelle markers affecting distinct cellular compartments. Typically, the efficacy of photodynamic therapy faces challenges in hypoxic conditions due to the dependence on oxygen molecules for ROS generation (*33–35*). However, our study unveils an escalated phototoxic impact of Mito-Tracker in hypoxia, driven by mechanisms independent of ROS but mitochondrial functions. Further investigations would lead to the development of more advanced photosensitizers specifically designed to target cancer cells resilient to hypoxic conditions.

## Discussion

A sensitive and quantitative way to assess phototoxicity in live cells holds significant importance for biologists seeking to comprehend the impact of drugs or fluorescent dyes on cell functions. Conventional assays, reliant on ensemble measurements or microscopy techniques detecting cell morphological changes, primarily evaluate late-stage phototoxicity. In this study, by using EB3-EGFP signals to quantify microtubule polymerization dynamics, we introduce a sensitive and quantitative method to measure early-stage phototoxicity and cellular perturbations. The dynamics of microtubule polymerization reflect cellular energy flux, which changes rapidly in response to cellular disturbance and phototoxicity. We studied the influences of different lasers and common fluorescent organelle markers on microtubule polymerizations within live cells. The findings suggest that distinct organelle markers exhibit varying phototoxic effects when exposed to different laser wavelengths. The utilization of the RPOC technology enables the comparison of interactions between lasers and dyes, whether colocalized or offset, leading to markedly distinct impacts on cell responses. Furthermore, a study of cell responses in hypoxia concludes that while no elevation is observed in the phototoxicity of ER-Tracker, the phototoxicity of Mito-Tracker is increased. The increased phototoxicity of Mito-Tracker in hypoxia does not appear to be associated with ROS generation, suggesting the involvement of alternative mitochondria-specific mechanisms contributing to cellular damage.

The developed method for quantifying EB3 comet dynamics and assessing phototoxicity is broadly applicable across diverse conditions. Comparative statistical analysis of EB3 comets, using images obtained from both a commercial confocal fluorescence microscope and our in-house confocal system, produced highly consistent results. Since circularity is unitless, the quantification of EB3 comets remains independent of image size, provided the comets are visible and generally spatially separated. Typically, an imaging speed exceeding 5 seconds per frame would yield comparable outcomes. The transfection of EGFP to EB3 proteins in other cell lines further extends the applicability of this method. Similarly, utilizing fluorescent protein-conjugated EB1 can also facilitate phototoxicity analysis (*36, 37*). Beyond the organelle markers employed in this study, this methodology enables the evaluation of the photosensitization effects of other fluorescent markers or drugs on cells. It can also be applied to comprehend the phototoxic effects induced by pulsed lasers utilized in nonlinear optical imaging modalities, both for labeled and unlabeled cells. When combined with RPOC, which enables site-specific light interactions with organelles, this approach allows for establishing improved correlations between organelle functions and phototoxicities.

## Methods and materials

### Microscopy and RPOC

A commercial confocal fluorescence microscope (LSM510, Zeiss) was applied for a portion of the fluorescence imaging in this study. The microscope is equipped with 7 laser lines including 405, 458, 477, 488, 514, 543, and 633 nm, and two photomultiplier tube (PMT) channels. For capturing images of EB3-EGFP comets, a 10 µW 488 nm laser was used in combination with a bandpass 505-530 filter. The images of 112.3×112.3 µm dimensions consist 512×512 pixels were acquired with a pixel dwell time of 1.61 µs. A total of 30 frames of images were acquired for each time-lapse image stack. The fluorescence signals from the H2DCFDA ROS sensor were acquired using the bandpass 505-530 filter. In **Figure 2**, imaging utilizes a 10 µW 543 nm laser and the maximum available power of the 633 nm laser (1.7 µW). A Zeiss 40X/1.3 oil objective lens was used for the acquisition of all images with the commercial confocal fluorescence microscope.

A lab-built confocal and RPOC system has four lasers including 405, 473, 532, and 589 nm. The workflow of RPOC is illustrated in **Figure S3**. For the excitation of the EB3-EGFP signals, a 473 nm laser at 20 µW is used, while the visualization of ER-Tracker signals employs a 589 nm laser at 60 µW. The EB3-EGFP signals are acquired by a PMT (H7422-40, Hamamatsu) with a bandpass filter (FF01-509/22, Semrock). Meanwhile, fluorescence signals from chemical labels are acquired in a separate PMT with a different bandpass filter (ET642/80m, Chroma Technology Corporation). RPOC enables precise generation of phototoxicity by activating a 532 nm laser with 8 µW at specified locations. The selection of APXs is achieved by using a comparator circuit box that allows for manually adjusting intensity thresholds. The RPOC system, alongside its ’invert’ function, allows for both the selection and deselection of APXs specifically on mitochondria based on the Mito-Tracker Red signals (*28*). RPOC operations utilizing the 532 nm laser involve 30 frames, totaling approximately 152 seconds, with a galvo mirror scanning speed set at 20 µs. An Olympus 60X/1.2 water objective lens is used for imaging and RPOC. Further details of the continuous-wave (CW) RPOC system are available in a previous report (*28*).

Stage top incubators from Tokai Hit (model OTH-STXF-WSKMX-CO2O2) are integrated into both the commercial confocal microscope and the RPOC system, allowing for the maintenance of both normoxic and hypoxic conditions (**Figure S4**). Cells are exposed to various durations in hypoxia before imaging and laser interactions. Within the hypoxic condition, the oxygen level is maintained at 0.1% alongside 5% CO_2_. The temperature within the culture dish is set to 37 °C.

### Quantitative image analysis

Images containing EB3-EGFP signals are processed using ImageJ. For the data obtained with the commercial confocal fluorescence microscope, time-lapse images containing tubulin dynamics comprising 30 frames are acquired both before and after treatment. The averaged image of the last 5 frames (26-30 frames) of the pre-treatment time-lapse represents the pre-treatment conditions, while the average of the initial 5 frames (1-5 frames) of the post-treatment time-lapse signifies the post-treatment conditions. For the data collected with the lab-built confocal fluorescence and RPOC system, a single timelapse image stack is acquired during the treatment phase. Here, the average of the first 5 frames (1-5 frames) represents the pre-treatment conditions, and the average of the last 5 frames (26-30 frames) represents the post-treatment scenarios. Processed tubulin-EGFP images are utilized for quantifying the circularity of the EB3 comets. The Candle-J denoising, an ImageJ plug-in, is employed to enhance the image quality and contrast of EB3 comets (*26*). The method for circularity quantification of EB3 comets is detailed in **Figure 1A**. A Gaussian-blur filter is applied with a sigma(radius) value set as 2. The Candle-J denoising uses default parameters from the ImageJ plugin. Circularity histograms are generated using the ImageJ histogram function and plotted using Origin 2021. Heatmaps displaying comet circularity below 0.6 are plotted using Origin 2021.

### Statistics

The quantitative analysis of H2DCFDA intensity in **Figure 5C** involves analyzing 11-12 cells.

### Cell preparation

HeLa Kyoto EB3-EGFP cells were purchased from Biohippo for the investigation of microtubule polymerization and EB3 dynamics. HeLa cells (no EGFP transfections) were obtained from ATCC to study ROS generation under various conditions. Cells were cultured in Dulbecco’s Modified Eagle Medium (DMEM, ATCC) with 10% fetal bovine serum (FBS, ATCC) and 1% penicillin/streptomycin (Thermofisher Scientific). The cells were plated in glass-bottom dishes (MatTek Life Sciences) with 2 mL culture medium and then incubated in a CO_2_ incubator at 37 °C and 5% CO_2_ concentration. The cells were allowed to grow to approximately 50-70% confluency and were directly used for live-cell imaging.

### Fluorescent labeling of ER, mitochondria, and nucleus

HeLa Kyoto EB3-EGFP cells were first seeded in glass-bottom dishes and allowed to culture overnight until reaching a confluency level of about 50-70%. All the organelle labeling fluorescent dyes were used following the recommended guidelines and protocols provided by the manufacturers. ER Tracker was introduced to the culture medium at a final concentration of 1 µM. Mito-Tracker Red CMXRos was introduced to the culture medium at a final concentration of 200 nM. Nuclei were stained by SYTO 82 Orange Fluorescent Nucleic Acid Stain at a final concentration of 0.5 µM. Subsequently, the cells were incubated for 30 minutes at 37 °C and 5% CO_2_ concentration before imaging. For mitochondria labeling and measurements involving CW-RPOC, cells were rinsed once with a warm culture medium subsequent to the incubation period with Mito-Tracker Red. The absorption spectra of ER-Tracker and Mito-Tracker were measured using a UV-VIS spectrometer (GENESYS 50, Thermofisher).

### ROS measurements using H2DCFDA

HeLa cells (without EGFP transfections) were first seeded in glass-bottom dishes and cultured overnight until they achieve a confluency of around 50-70%. Subsequently, H2DCFDA was added to the culture medium at a final concentration of 10 µM, while ER Tracker Red or Mito-Tracker Red CMXRos was introduced at concentrations of 1 µM or 200 nM, respectively. The stock solution of H2DCFDA was prepared inside a glove box with a concentration of 20 µM. The prepared stock solution was utilized within a limited timeframe to prevent degradation. The necessary fluorescent dyes were then added to the cells, and the cells were incubated for 30 minutes at 37 °C under a 5% CO_2_ concentration before the imaging.

## Acknowledgment

This research is supported by NIH R35GM147092 and a pilot grant from the Purdue Center for Cancer Research awarded to C. Zhang. We also thank Mr. Wen Xiu from Dr. Chris Uyeda’s group for assisting us with the handling of H2DCFDA in their glove box.

## Author contributions

C. Zhang and S. Mahapatra designed the experiments. S. Mahapatra performed data acquisition and analysis. S. Ma assisted in sample preparation and data acquisition. B. Dong assisted in RPOC instrumentation and data acqusition. C. Zhang and S. Mahapatra wrote the manuscript.

