## Supplementary Information for "Quantification of cellular phototoxicity of organelle stains by the dynamics of microtubule polymerization"

### Supplementary notes

#### Rapid disruption of microtubule dynamics facilitated by laser exposure to the Nucleic Acid Stain

We investigated the phototoxic impact of SYTO Orange Nucleic Acid Stain on HeLa Kyoto EB3-EGFP cells. Upon exposure to only 10 µW 488 nm laser scanning, we observed rapid disruption of microtubule dynamics, evidenced by the diffused EGFP signal in the initial frames as shown in **Figure S1**. This impairment can be attributed to the phototoxic properties of the nucleic acid stain. Cytotoxic effects are unlikely since the cells retained vitality with healthy microtubule dynamics even after a 30-minute incubation period. Furthermore, upon zooming out, untreated cells exhibit normal microtubule dynamics. The observed toxicity stems from the combined impact of the excitation laser and the nucleic acid stain. Note that the 488 nm laser used here is not the optimal wavelength for the excitation of the Nuclei Acid Stain but for the EGFP.

#### Phototoxicity of Mito-Tracker Red in hypoxia

The Mito-Tracker Red displays a broad absorption spectrum, as shown in **Figure S2**. Its absorption peak is approximately 590 nm, similar to that of the ER-Tracker Red (**Figure 2E**). Among available laser options in our commercial confocal fluorescence microscope, the 543 nm laser best matches the excitation needs of these dyes. Our investigations into normoxia and hypoxia involved the use of the 543 nm laser to examine cells labeled with Mito-Tracker and cultured within the stage-top incubator (**Figure S4**). As hypoxia duration increased, EGFP signals decreased due to oxygen deficiency. Furthermore, HeLa cells labeled with Mito-Tracker and illuminated with the 543 nm laser showed elevated susceptibility to increased hypoxic conditions, evidenced by the substantial membrane blebbing as shown in **Figures 5A and S5A**. Using normal HeLa cells treated by the ROS sensor H2DCFDA, we quantified the level of ROS generated under various treatment conditions. Hypoxia exposure significantly reduced the generation of ROS after laser illumination of HeLa cells labeled with MitoTracker (**Figures 5B,C, and S5B**). This suggests that the elevated phototoxicity observed in hypoxia when exciting Mito-Tracker with the 543 nm laser is not due to ROS generation but possibly involves other mechanisms, which will be explored in future studies.

**Phototoxicity of ER-Tracker Red in hypoxia**

Under normoxic conditions, when subjected to the same laser wavelength, ER-Tracker Red displayed comparatively less phototoxicity in contrast to Mito-Tracker. To assess the phototoxic effects more effectively, we utilized the 589 nm laser on a lab-built confocal fluorescence microscope for treatment purposes. Following a 152-second laser exposure, only a minor decline in microtubule polymerization was noted in normoxic conditions. Hypoxic exposure did not substantially increase the microtubule fragmentation as shown in **Figure S6A**. Employing a ROS sensor H2DCFDA, we detected a significant increase in ROS generation after 589 nm illumination of the ER-Tracker treated cells. This increase occurred more rapidly in the normoxic condition compared to the hypoxic condition (**Figure 6C**), suggesting a reduction in ROS generation in hypoxia. Subsequently, we subjected the labeled cells to a much longer 20-minute treatment using the 589 nm laser. Despite this treatment resulting in complete disruptions in microtubule polymerization in both cases (**Figure S7**), no membrane blebbing was observed in the hypoxia condition, unlike what was seen in the case of Mito-Tracker (**Figure 5A**).

**Supplementary figures**

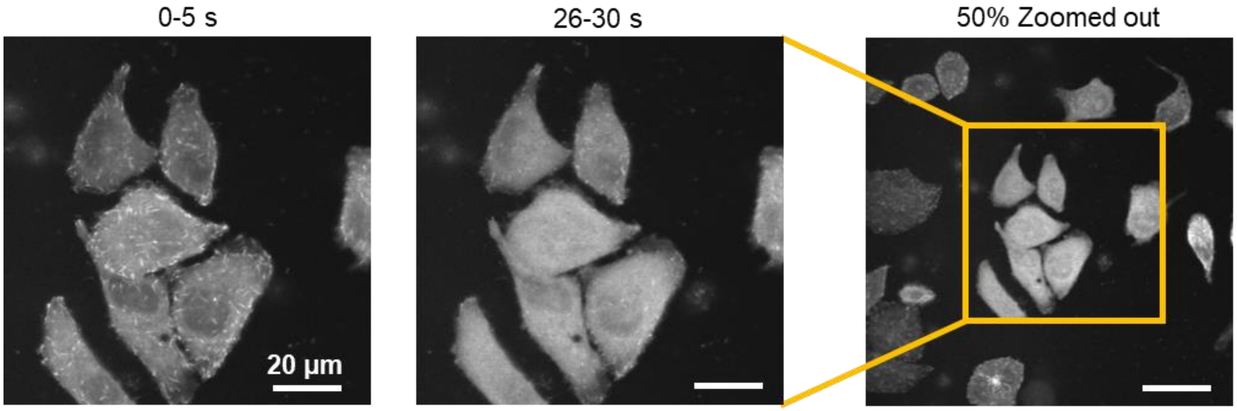


**Figure S1.** Fluorescent images of EB3-EGFP signals within HeLa cells, illuminated by a 488 nm laser via laser scanning. The cells are treated with SYTO 82 Orange Fluorescent Nucleic Acid Stain at a final concentration of 0.5 µM.

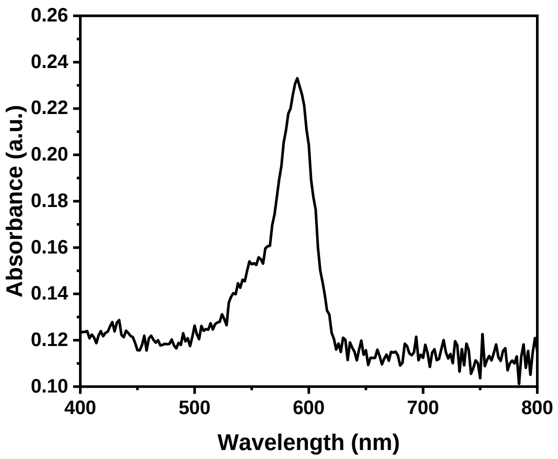


**Figure S2.** The absorption spectrum of Mito-Tracker Red CMXRos.

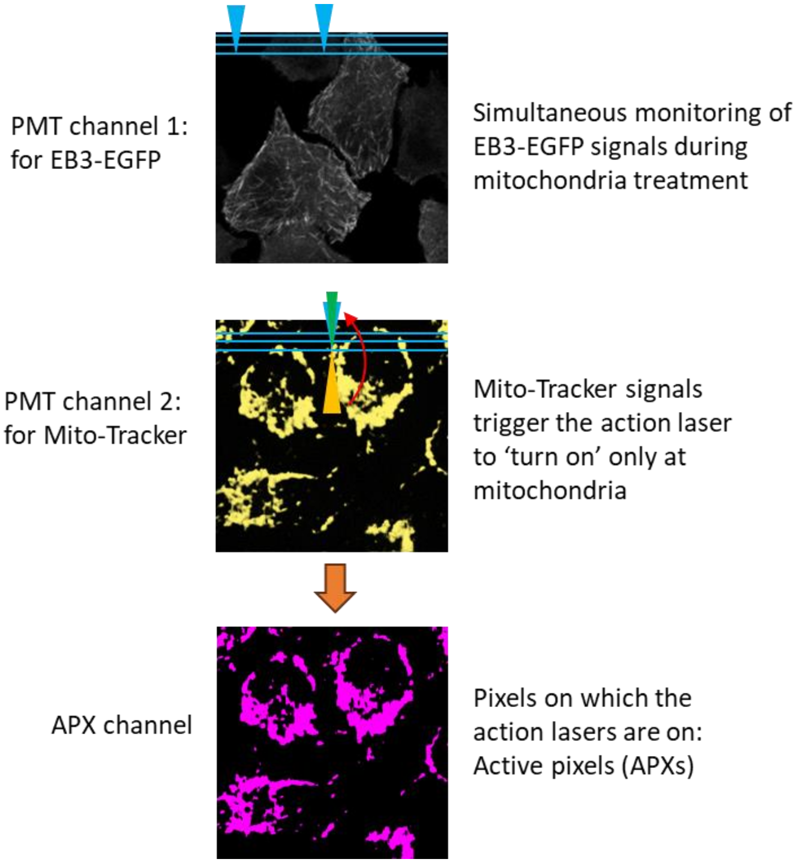


**Figure S3**. The workflow of real-time precision opto-control (RPOC). In the figures, blue arrows represent the excitation laser, yellow denotes the optical signal from targets, and green signifies the action laser. PMT refers to the photomultiplier tube, while APX stands for active pixels.

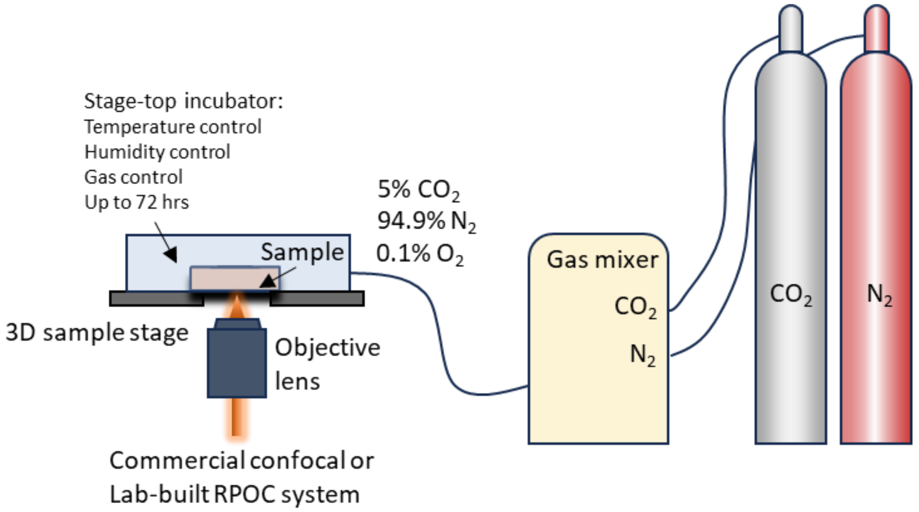


**Figure S4.** The stage-top incubation system connected to gas cylinders and a gas mixture to establish both hypoxia and normoxia culture conditions.

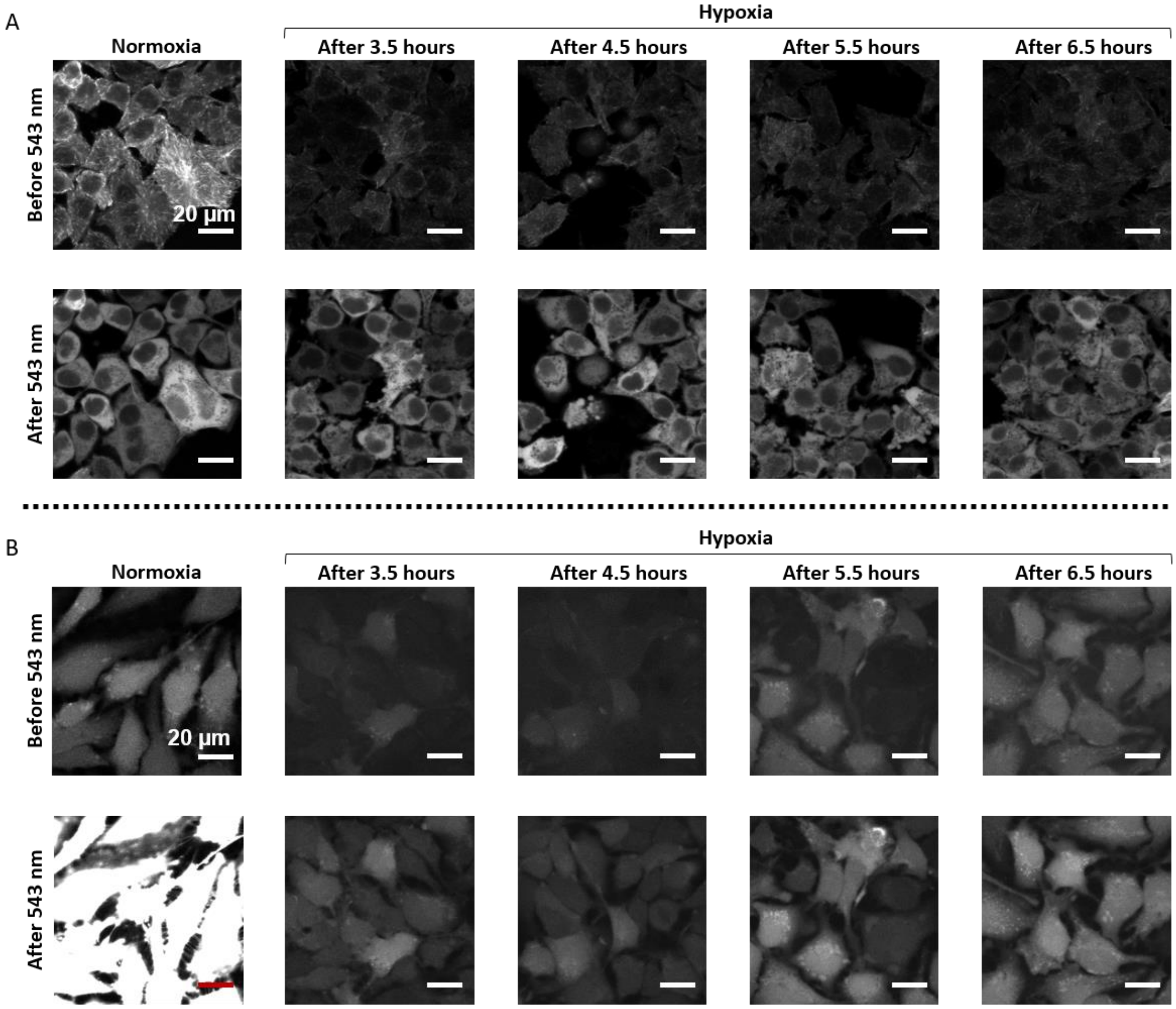


**Figure S5.** (A) Fluorescent images depicting EB3-EGFP signals in HeLa cells illuminated by the 543 nm laser for 20 minutes under normoxic and various hypoxic exposure durations. The hypoxic conditions additionally trigger significant membrane blebbing that indicates elevated phototoxicities. (B) Evaluation of ROS generation induced by MitoTracker before and after the 543 nm laser illumination under normoxic and various hypoxic exposure durations. The fluorescent signals originate from HeLa cells incubated with H2DCFDA. A 10 µW 488 nm laser is used to excite the H2DCFDA signals.

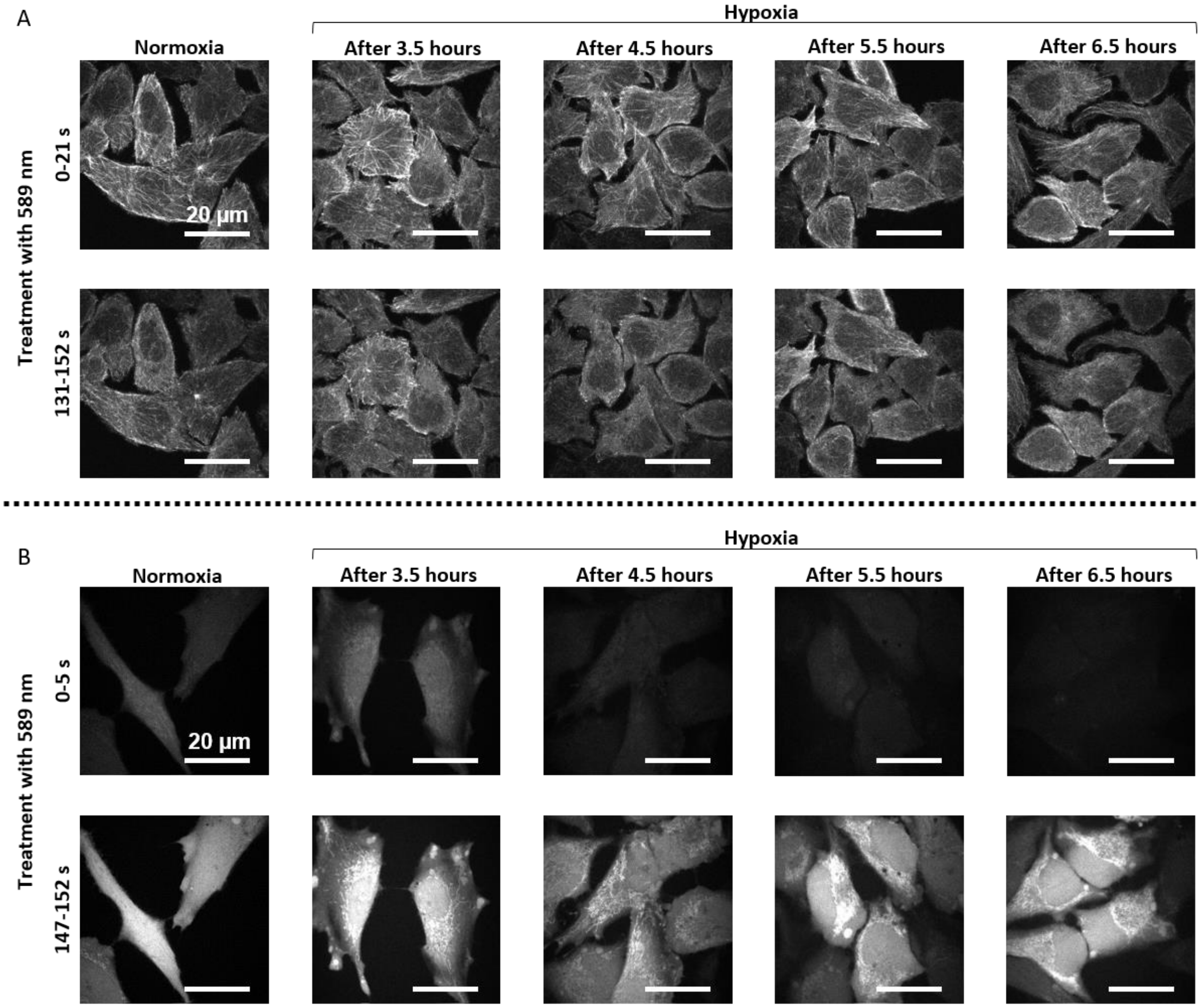


**Figure S6.** (A) Fluorescent images depicting EB3-EGFP signals in HeLa cells illuminated by the 589 nm laser for 152 seconds under normoxic and various hypoxic exposure durations. Additionally, a 20 µW 473 nm laser is concurrently scanned with the treatment lasers to excite the EGFP signals. (B) Evaluation of ROS generation induced by ER-Tracker before and after the 589 nm laser illumination under normoxic and various hypoxic exposure durations. The fluorescent signals originate from HeLa cells incubated with H2DCFDA. A 20 µW 473 nm laser is used to excite the H2DCFDA signals.

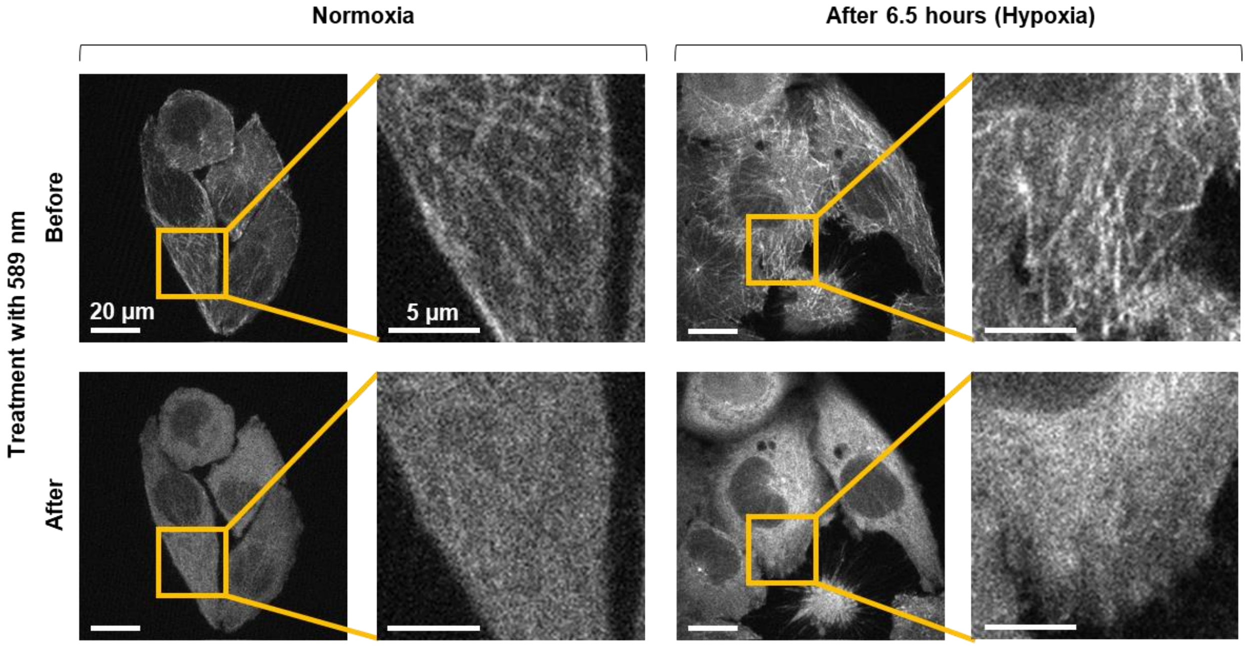


**Figure S7.** Fluorescent images depicting EB3-EGFP signals in HeLa cells illuminated by the 589 nm laser for 20 minutes under normoxic and 6.5 hr hypoxic exposure. Specific regions within the images are magnified to enhance the display of cellular details. No membrane blebbing is observed in either case.

**Supplementary Video 1.** The dynamics of EB3 and microtubule polymerization in HeLa cells revealed by fluorescent microscopy.

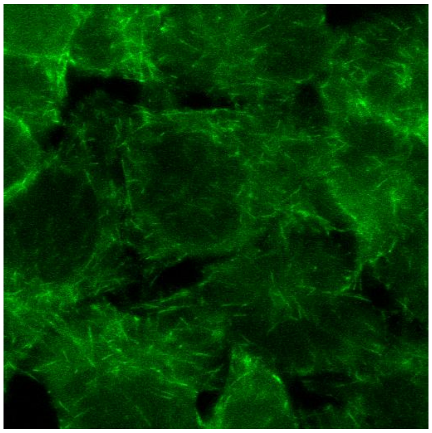
