## Supplementary figures and images for "Quantification of cellular phototoxicity of organelle stains by the dynamics of microtubule polymerization"

### Figure S1

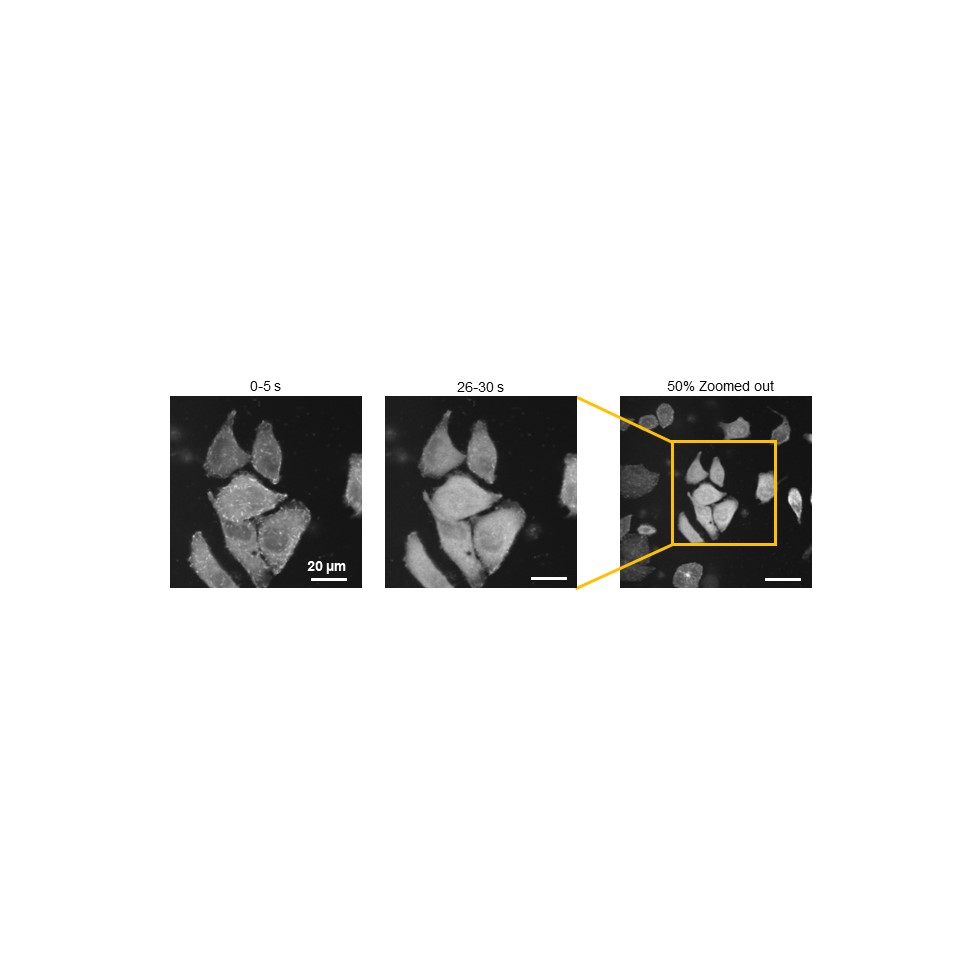

### Figure S2

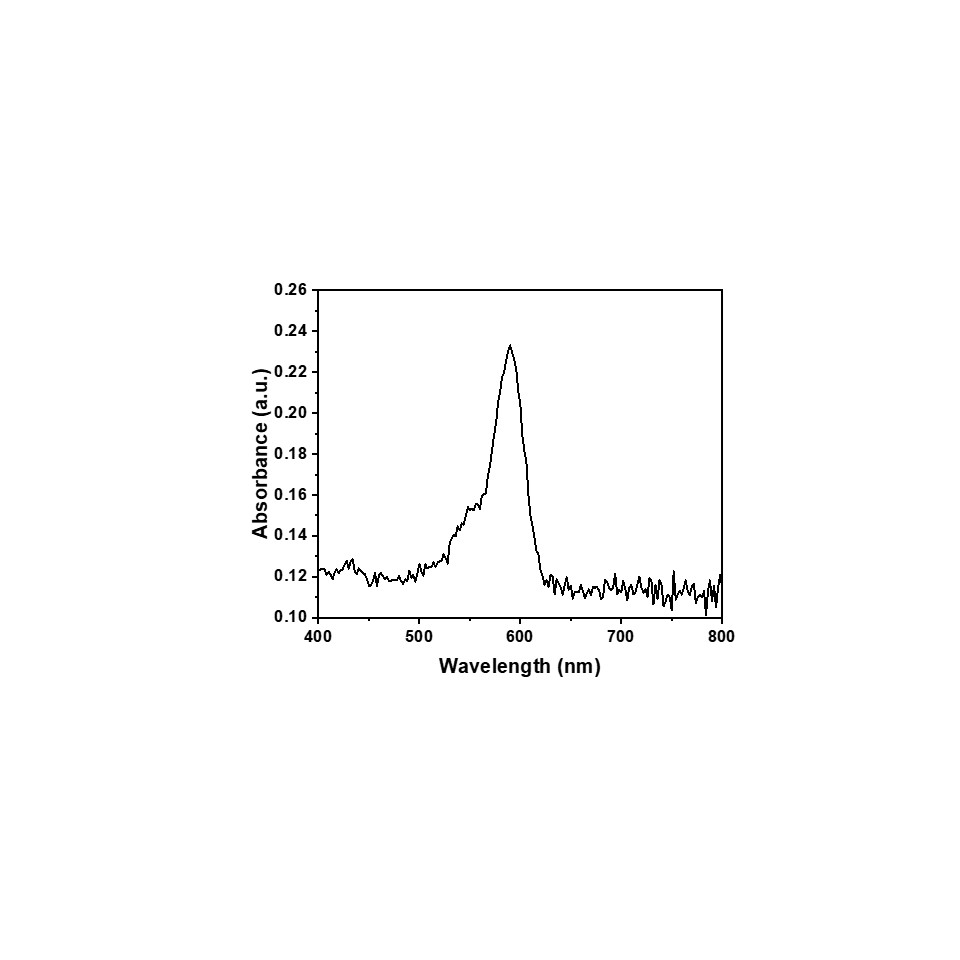

### Figure S3

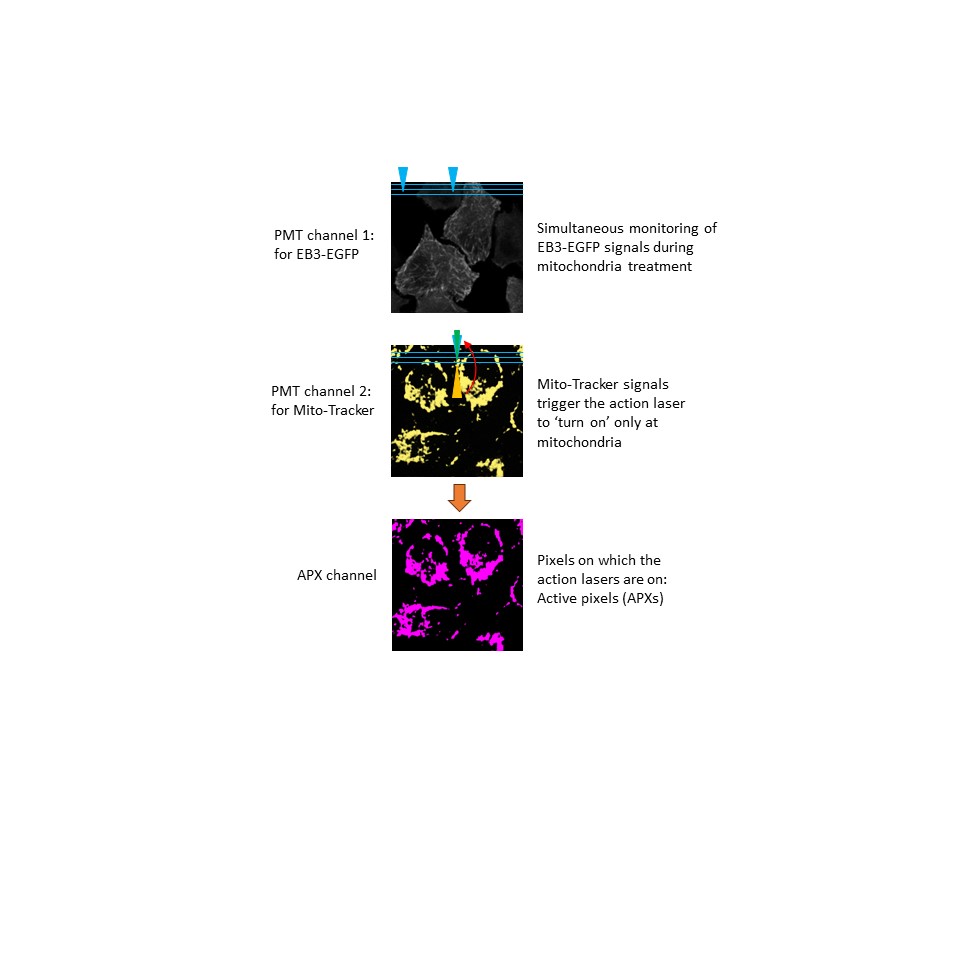

### Figure S4

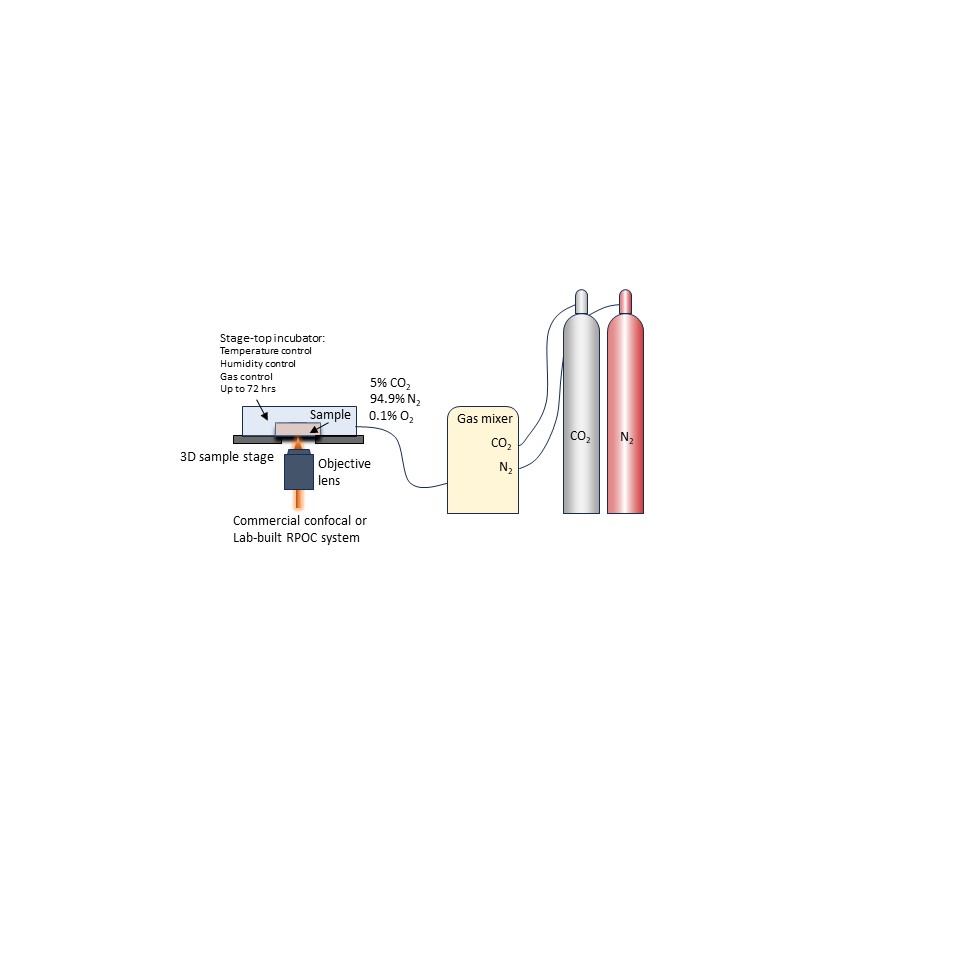

### Figure S5

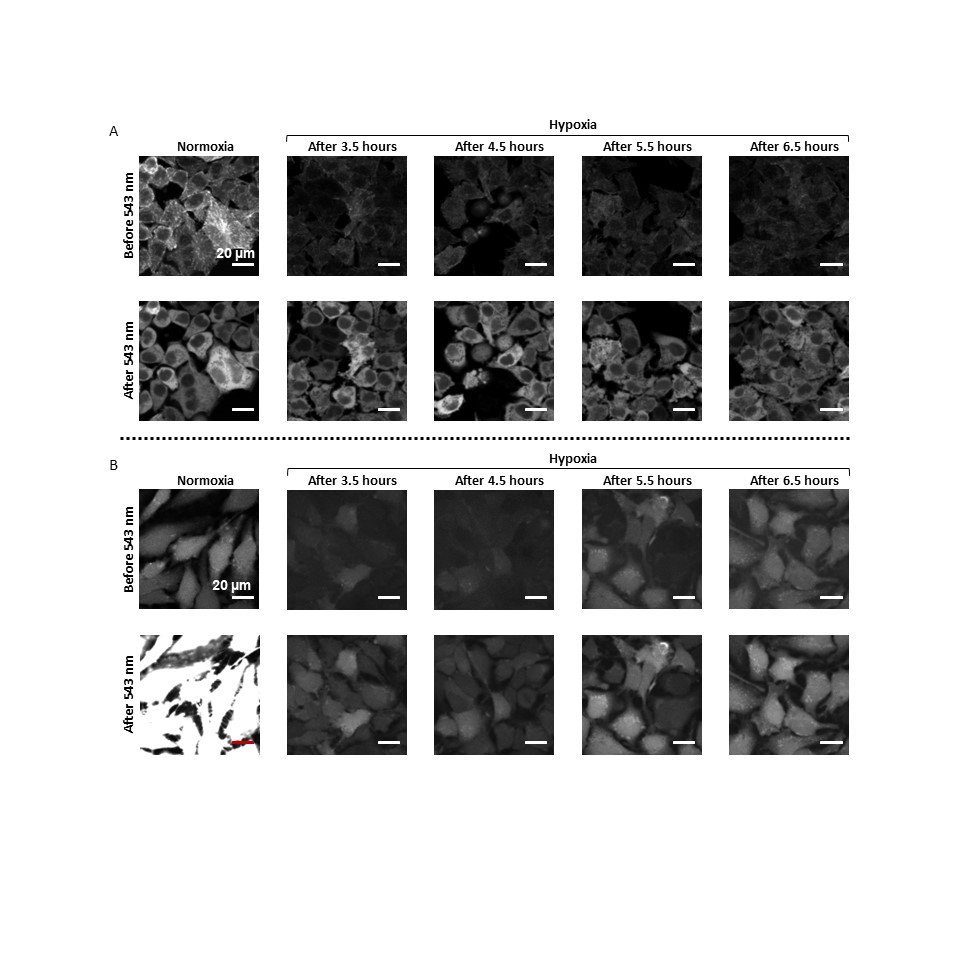

### Figure S6

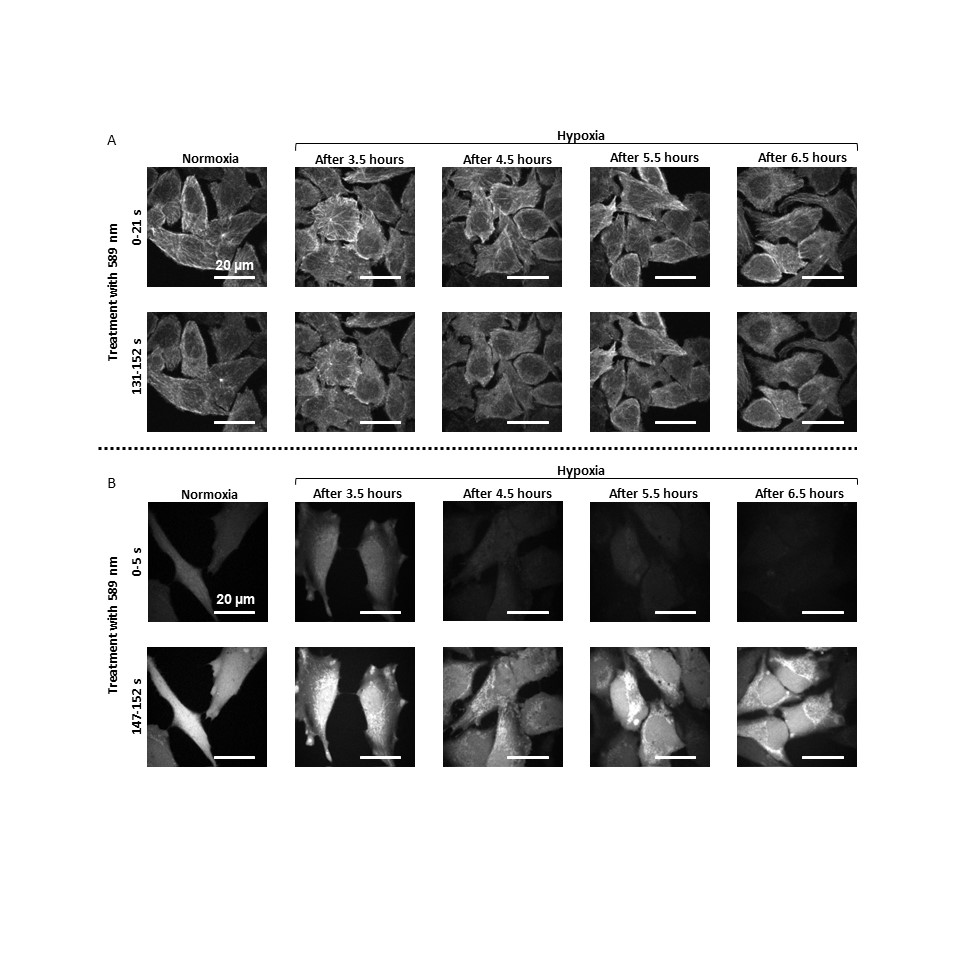

### Figure S7

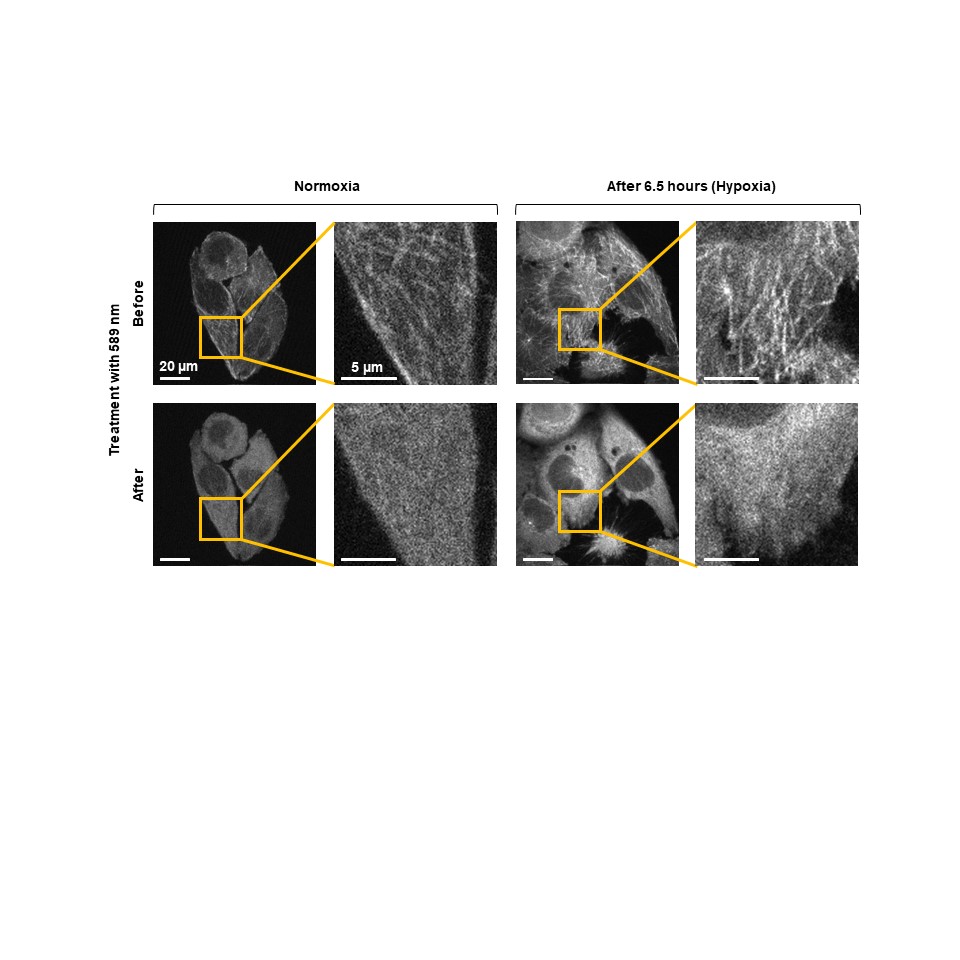
